# Evolution of lower levels of inter-locus sexual conflict in *D. melanogaster* populations under strong selection for rapid development

**DOI:** 10.1101/2021.02.08.430125

**Authors:** Avani Mital, Manaswini Sarangi, Snigdhadip Dey, Amitabh Joshi

## Abstract

*D. melanogaster* laboratory populations subjected to selection for rapid development and early reproduction have been found to have evolved reduced adult body size and lower levels of inter-locus sexual conflict compared to their ancestral controls. We tested the contribution of a smaller body to the evolution of reduced sexual conflict in these populations, since body size differences are known to affect sexual conflict levels in this species. We cultured larvae from the control populations at high density to obtain flies as small as those from the selected populations. The effect of body size reduction on sexual conflict was asymmetric, with smaller body size resulting in reduced male manipulative ability but not female resistance to mating-induced harm. These results were not due to differences in behavioural patterns of smaller flies, such as differences in overall mating exposure of females to different types of males. We hypothesize that evolution for rapid development and the correlated reduction in body size has resulted in lower male manipulative ability, and sexually antagonistic co-evolution has lowered female resistance to such manipulations. These populations have also evolved incipient reproductive isolation from their controls, likely through sexual conflict (reported earlier), and our results support the view that this is an outcome of strong, directional selection for rapid development.

## Introduction

Theory suggests that, in general, shorter development times can yield consistent fitness benefits by reducing the probability of death till fitness is realized, and by increasing generational turnover rate (Roff 2002). However, time spent in development directly affects many aspects of the adult phenotype and, thus, can have major consequences for fitness. The fitness impact of development time is particularly large in holometabolous insects as they acquire resources mainly during pre-adult development. Experimental evolution studies on populations of *Drosophila melanogaster* have demonstrated clear trade-offs that have potentially limited the evolution of very short generation times in the wild (Zwaan et al. 1995, Nunney 1996, Chippindale et al. 1997, Prasad et al. 2000, 2001, Modak et al. 2009, Satish 2010), with many of these trade-offs manifesting via reduced adult resources available to the fly as a consequence of shortened larval development.

Reduced body size of adults can result in significant fitness loss through lower pre-adult survivorship (Chippindale et al. 1997, Prasad et al. 2000), and life-span (Prasad 2004). Sex specific differences in fitness loss include lower egg output (Stearns 1992, Lefranc and Bundgaard 2000) and reduced attractiveness towards males (Partridge et al. 1987a, Long et al. 2009, Lupold et al. 2011, Nandy et al. 2012) in smaller females. There are numerous studies linking male body size with mating ability, especially under competitive conditions. For example, mating and re-mating frequency, copulation duration, courtship vigour (Partridge and Farquhar 1983, Partridge et al. 1987a, b, Markow 1988, Markow and Ricker 1992, Lefranc and Bundgaard 2000, Friberg and Arnqvist 2003, Wigby et al. 2016) are lower for relatively smaller males, as is total investment in seminal fluid proteins that aid in sexual competition (Bangham et al. 2002, Pitnick and Garcia-Gonzales 2002, but see Wigby et al. 2015). The amount of resources available to adults can, therefore, constrain the reproductive success of both sexes.

Males and females exhibit different reproductive strategies to maximize fitness: male fitness increases with increasing number of mates, female fitness with number of eggs laid. This creates the selective environment for one sex to modify the behaviour and physiology of its partner, in order to reap maximum benefit from a mating (Bateman 1948, Trivers 1972, Parker 1979) which, in turn, gives rise to inter-locus sexual conflict (Holland and Rice 1998, Chapman et al. 2003, Arnqvist and Rowe 2005). In *D. melanogaster*, males produce various proteins in their accessory glands (Acps), which mediate such conflict, and are transferred to the female in the ejaculate (Chapman et al. 1995, Rice 2000). Subsequently, females can respond to such manipulations through the evolution of resistance, resulting in a co-evolutionary arms race between the sexes (Chapman et al. 2003, Arnqvist and Rowe 2005).

Since smaller adults have limited resources to invest in reproduction related traits, small males are likely to be less harmful, and small females less resistant, than their normal sized counterparts. Thus, selection for rapid development (Chippindale et al. 1997, Prasad et al. 2000) can reduce levels of inter-locus sexual conflict via body size reduction. This has been suspected in the past in populations of *D. melanogaster* selected for rapid development and early reproduction in our laboratory (M Ghosh and Joshi 2012). Females from the selected populations showed higher mortality rates than females from control populations when they mate with males from control populations, and this increased mortality also resulted in incipient reproductive isolation between the selected and control populations. Furthermore, micro-array studies on the selected populations have also revealed significant down-regulation of various Acps in young males (Satish 2010).

In this study, we have used the four replicate faster developing and early reproducing *D. melanogaster* populations (shorter generation times), and their ancestral controls, to address the following question: how body size reduction (as a consequence of selection for rapid development), affects a) mate manipulative ability of males, and b) female resistance to mate induced harm.

We assayed inter-locus sexual conflict in two ways: by estimating 1) post mating mortality of females to directly assess mating induced harm, and 2) male success in manipulation of female behaviour using females from an unrelated population. The control population flies are more than twice as large as those from the selected populations (M Ghosh and Joshi 2012), a substantial difference in body size that could have large effects on mate harming abilities of males. By limiting the food available to the larvae of the control populations for a single generation, we were able to collect adults that did not significantly differ in body size from those of selected populations, and, thus, distinguished between the effects of body size reduction and those of genetic changes that the selected populations would have undergone as a consequence of selection for rapid development and early reproduction.

## Materials and Methods

### Experimental Populations

The studies reported here were conducted on four matched pairs of *D. melanogaster* populations (control and selected) maintained under constant light conditions, ad libitum banana-jaggery food medium, relative humidity of around 80% and a large breeding number ranging from 1500-1800 per population. The maintenance protocol for the control populations has been described in detail earlier (Sheeba et al. 1998). Briefly, the control populations, JB (**J**oshi **B**aseline), experience a 21 day discrete generation cycle where cultures are maintained for the first 12 days in glass vials with approximately 6 mL food medium and around 60-80 eggs per culture. All eclosing flies from each culture are provided fresh food in vials every alternate day until 18 days from initiation of the cultures. They are subsequently collected in plexi-glass cages and provided a smear of live yeast paste on food medium for three days. Flies are then allowed to lay eggs for 18 hours before cultures for the next generation are started. The selected populations, FEJ (**F**aster-developing, **E**arly-reproducing, **J**B), first described in Prasad et al. (2000), are maintained in similar culture conditions in vials for 6 days. Subsequently, the cultures are checked every two hours for the first 15 or more flies that have emerged per culture vial, which are directly collected into a plexi-glass cage provided with food medium with live-yeast paste. Thus, selection for the fastest developing ~20% of flies from each culture is imposed at this stage. Three days later the flies are allowed to lay eggs for an hour before cultures for the next generation are initiated, thus imposing selection for early reproduction relative to the controls. Studies reported here were conducted between 510-540 generations of FEJ selection, corresponding to about 260-275 JB generations, the generation time of the FEJs having come down to ~10 days.

Each of the four FEJ_(1-4)_ populations has been derived from one JB_(1-4)_ population (matched subscripts) such that ancestrally, FEJ_*1*_ is more closely related to JB_*1*_ population than to other FEJ populations. Because ancestry is accounted for, selected and control populations that share the same numerical subscript have been treated as blocks in the statistical analyses. Different populations of the same block were assayed together.

Since the selected and control populations experience different maintenance regimes, non-genetic parental effects can be confounded with traits that have evolved in different populations. This was controlled for by rearing both control and selected populations under similar conditions for at least one full generation (*standardized* populations) before assaying any trait. This involved collecting all eclosing flies in a plexi-glass cage and providing the same time duration as controls to breed before eggs were collected from these standardized populations for various assays.

### Obtaining body size control flies

The experiments described here were performed with three sets of flies: the FEJ, JB and a body size control created for JB flies (Jc) by limiting food medium for the larvae for one generation to a sufficient degree such that the adults from these cultures were as small as FEJ flies. This was checked by measuring their dry weight which did not significantly differ from the dry weight of FEJ flies, as measured at the age at which these flies were used in the assay (Supplementary Material). For obtaining small JB flies (hereafter ‘Jc’), a larval density of 220-230 eggs /1.5 mL of banana-jaggery food medium was used, whereas for FEJ and JB flies, egg collection was done as for regular population maintenance.

### Collecting adults for assays

Since all experiments assayed adult behaviour, and FEJ and JB flies have different development times, collection of eggs from the populations was staggered so as to obtain adults of the same age on the same day. Prior to assaying, eggs from standardized JB and FEJ populations were collected and then freshly eclosed flies were separated as virgins within 6 hours from eclosion using light carbon dioxide anesthetization. Virgin males and females were subsequently housed in single sex vials and were aged for 3-4 days until the assays were performed.

### Post mating female mortality (50 hours of exposure to males)

We assayed the post-mating mortality of the three types of females under study (FEJ, JB and Jc) when paired with FEJ, JB or Jc males as mates. Pairs of flies were set up in vials with food, with 10 vials for each of the nine combinations, using light carbon dioxide anesthetization, and housed for 50 hours after which the males were removed from the vials without anesthesia. Females were thereafter kept individually in vials to assess mortality by checking for female death once every 24 hours. They were shifted to fresh food vials every alternate day for 14 days, at the end of which the cumulative mortality of the females was calculated.

### Refractory period assay

The purpose of performing this assay was to assess two components of mate manipulative ability of males, namely the ability to re-mate with an already mated female, and the ability to keep a female non-receptive to a second male after mating. For this purpose, a common female background was chosen to avoid confounding effects of using females from any of the populations being studied, as evolved males from that population may have an advantage in reproductive interactions, having co-evolved with females from the same populations for many generations. Scarlet eye (SE) mutant outbred populations that have been maintained in the laboratory for about 70 generations were used for this purpose. These populations were created by introgressing the SE gene into an MGB background. The MGB populations were created by mixing the four replicate JB populations and then re-deriving five new outbreeding populations from the resulting four-way hybrid populations.

For assessing the ability to mate with an already mated female, one virgin male and one virgin female each from the SE population were placed in vials using light anesthetization with carbon dioxide. Immediately after the first successful mating (minimum of three minutes in copula) was achieved, the male was replaced (without using anesthesia) with a virgin experimental male (FEJ, JB or Jc). Initially, 15 such vials were set up for each of the three regimes, and second mates were introduced into vials in which the first mating happened within four hours from setup. Two indices were used to assess the ability of JB, FEJ and Jc males to mate with a previously mated female: (1) the proportion of vials in which a second successful mating was *not* achieved (refractoriness), and (2) the time taken from introduction of the second mate till successful mating (refractory period). The ability to secure a mating with non-virgin female can involve making the female receptive for a second mating through forced copulation /coercion or successful courtship. In this assay, however, it was not possible to differentiate between these two mechanisms of second-male mating success.

To assess the ability of JB, FEJ and Jc males to keep the female they mated with non-receptive to a second male, a similar assay was carried out except that the first mate for the virgin SE female was the experimental male (FEJ, JB or Jc), and the virgin SE male was used as the second mate. Fifteen such vials were initially set up using light carbon dioxide anesthetization. The second mate virgin SE males replaced the first males without anesthesia in vials in which a successful mating was achieved within six hours. As in the previous case, refractoriness and refractory period were estimated. For both assays, the total observation period for both mating events to occur was eight hours.

### Statistical analyses

For post mating female mortality assay, the arcsine square root transformed fraction of females that died by day 14 was used for statistical analyses. Mixed model analysis of variance (ANOVA) was performed on these means with selection regime and male and female type (FEJ, JB or Jc) as fixed factors for analysis of female mortality. Mean refractory period across females for each block was subjected to mixed model ANOVA with male type as a fixed factor. Refractoriness data were also fractional data and therefore also subjected to arcsine square root transformation and then subjected to a mixed model ANOVA. Separate analyses were done for assays in which experimental males (JB, FEJ or Jc) were the first or second mate, respectively. For all analyses, means across replicate vials (population mean values) were used and FEJ, JB and Jc populations with the same numerical subscript were treated as random blocks for the ANOVA. All multiple pairwise comparisons were made by Tukey’s HSD test at a 0.05 significance level. All analyses were implemented on Statistica™ for Windows Release 5.0 B (StatSoft Inc., 1995).

## Results

### Post-mating female mortality (50 hours of mate exposure)

Cumulative female mortality through the first 14 days after exposure to males showed FEJ females mated with JB males to have substantially higher mortality than any other male-female type combination (Fig. 1). The ANOVA on cumulative mortality over 14 days revealed a significant main effect of female type, and multiple pair-wise comparisons revealed that, on average, FEJ females suffered significantly greater mortality than either JB or Jc females (Table 1, Fig. 2A). On an average, JB males induced substantially higher cumulative female mortality (Fig. 2B) with a significant main effect of male type (Table 1). There were also significant interactions between male and female type (Table 1) and multiple pair-wise comparisons revealed that FEJ females × JB males had significantly greater cumulative mortality than all other male-female type combinations at (Fig. 1) with the main effects of male type and female type are also likely to have been driven by this particular combination. Interestingly, FEJ female mortality was no greater when they mated with Jc males as compared to FEJ males, and Jc female mortality, similarly, was not different from JB female mortality when mated with JB, Jc or FEJ males indicating that Jc females, although smaller, tolerated mate induced harm as well as JB females but small Jc males showed reduced manipulative harm compared to JB males.

**Fig. 1.**
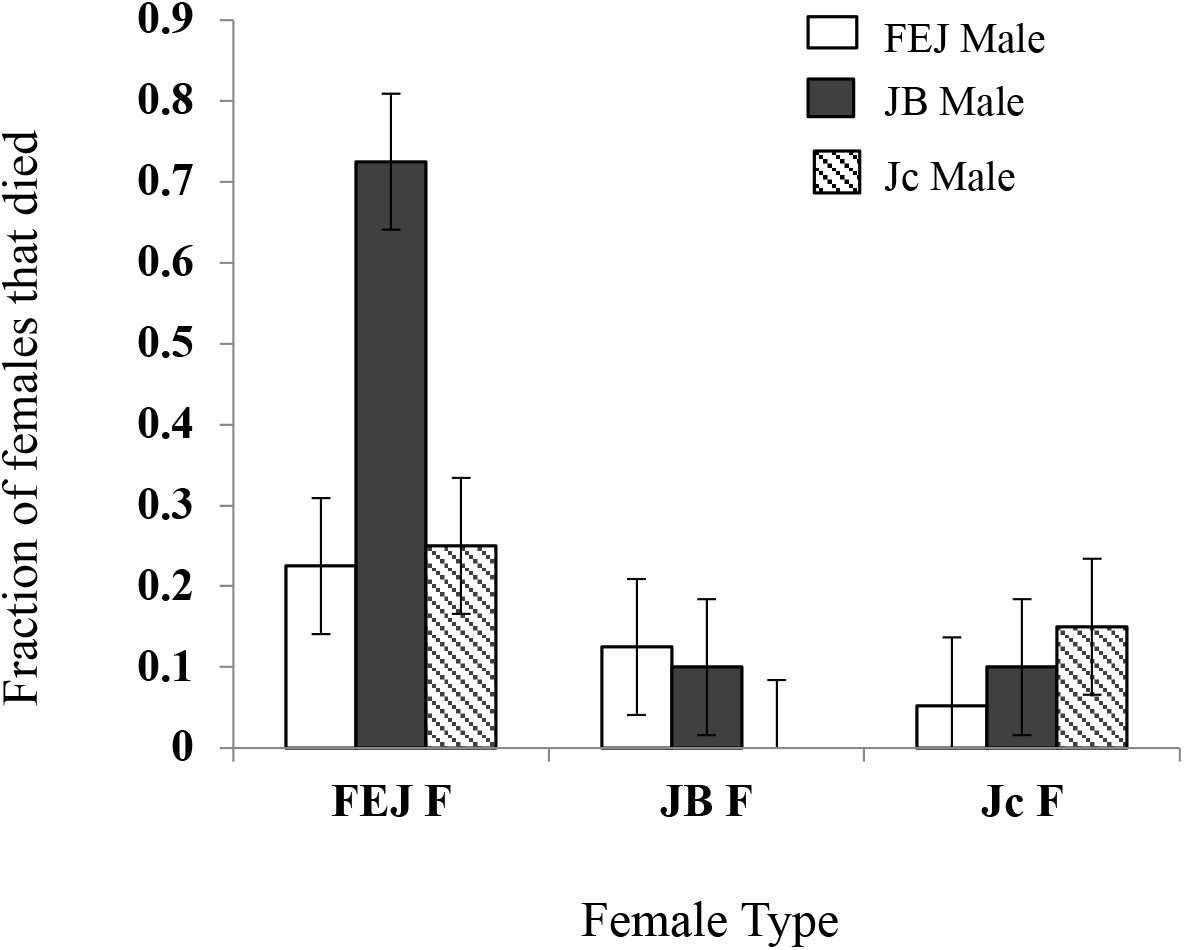
Mean fraction of females that died over the first 14 days from set up of the female mortality assay. Error bars are 95% confidence intervals allowing for visual hypothesis testing.

**Fig. 2.**
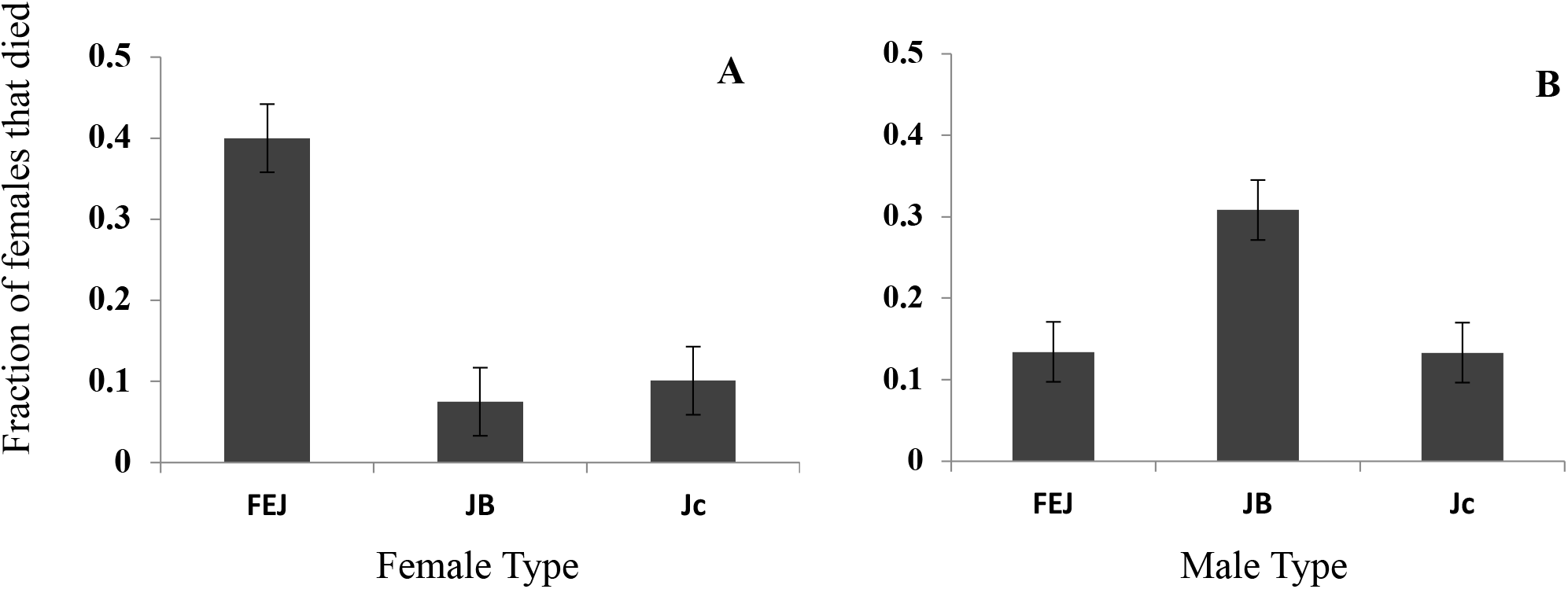
Mean fraction of females that died over the first 14 days from set up of the female mortality assay showing **A)** main effect of female, and **B)** main effect of male. Error bars are 95% confidence intervals, allowing for visual hypothesis testing.

**Table 1.** Results of ANOVA performed on the arcsine square root transformed cumulative female mortality by day 14, with male type and female type as fixed factors. In this design random factors and interactions are not tested for significance and have been omitted for brevity.

| Effect | *df* | MS | *F* | *P* |
| --- | --- | --- | --- | --- |
| Female | 2 | 0.8065 | 37.35 | 0.0041 |
| Male | 2 | 0.1726 | 6.976 | 0.0271 |
| Female × Male | 4 | 0.1954 | 5.594 | 0.0088 |

**Table 2.** Results of ANOVA performed on the refractoriness (fraction of SE females that did not re-mate) and refractory period (time taken for SE females to re-mate) when the first mate was the experimental male (FEJ, JB or Jc). In this design random factors and interactions are not tested for significance and have been omitted for brevity.

| Effect | *df* | MS | *F* | *P* |
| --- | --- | --- | --- | --- |
| Refractoriness | 2 | 0.1228 | 7.002 | 0.0269 |
| Refractory period | 2 | 2.3048 | 11.794 | 0.0083 |

**Table 3.** Results of ANOVA performed on the refractoriness (fraction of SE females that did not re-mate) and refractory time (time taken for SE females to re-mate) when the second mate was the experimental male (FEJ, JB or Jc). In this design random factors and interactions are not tested for significance and have been omitted for brevity.

| Effect | *df* | MS | *F* | *P* |
| --- | --- | --- | --- | --- |
| Refractoriness | 2 | 0.2962 | 43.631 | 0.0002 |
| Refractory period | 2 | 0.0593 | 0.0689 | 0.9341 |

### Refractory period assay

#### A) Ability to keep a mate non-receptive after mating

When the first mate was the experimental male (JB, FEJ or Jc), the ANOVA on mean refractory period showed a significant effect of male type (*F*_*2,6*_=11.79, *P=*0.008) (Fig 3A). JB males kept their mates non-receptive for a significantly longer time compared to the FEJ males, with Jc males being intermediate between FEJ and JB and significantly different from both (Tukey’s HSD at 0.05 significance level). Similarly, the fraction of females that did not re-mate (refractoriness) was significantly higher when the first mates were JB males, showing their greater ability to prevent mating of their mates compared with other males (*F*_*2,6*_=7.002, *P*=0.026) (Fig. 3B). However, the Jc males did not significantly differ from the FEJ males in the refractoriness they induced. Both these indices indicate a significant contribution of both genetic background and body size on the differences in mate manipulative ability between FEJ and JB populations.

**Fig. 3.**
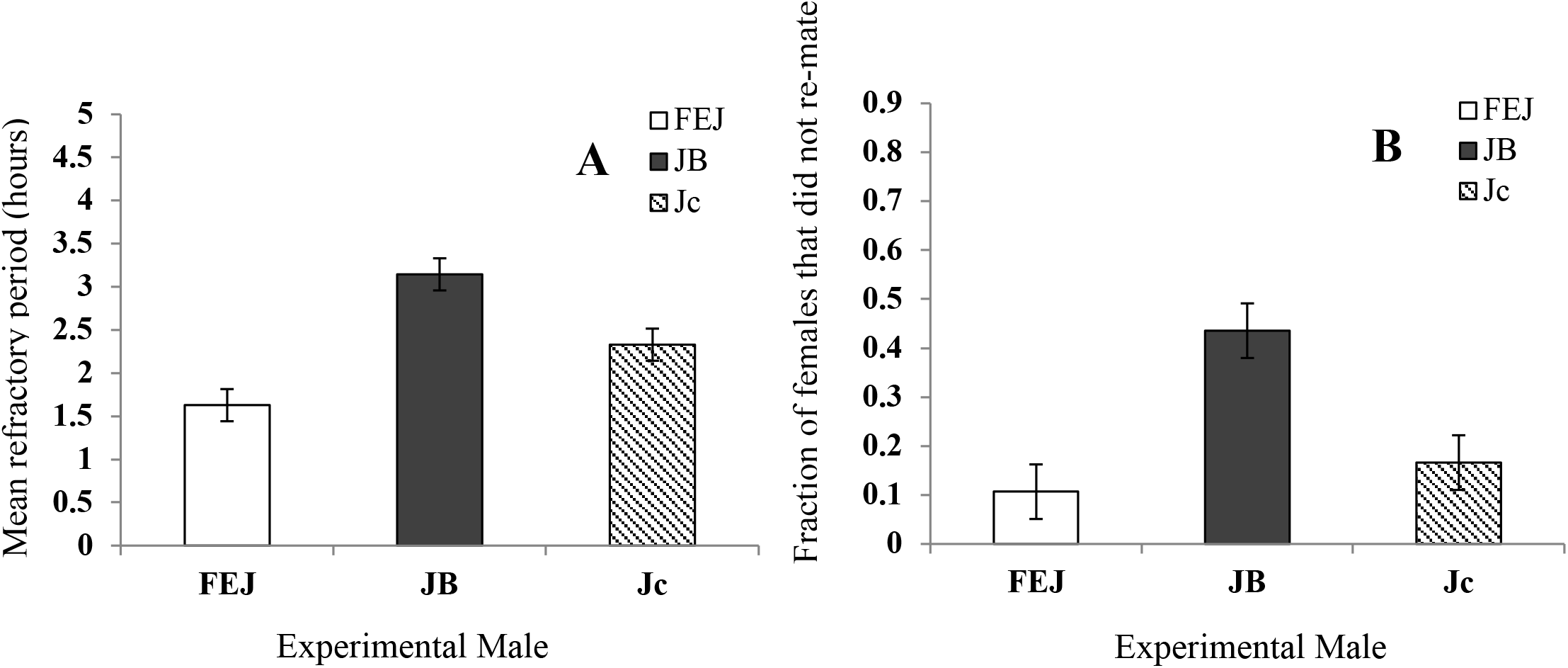
**A)** Mean refractory period: time till successful mating from introduction of second mates, first mates being FEJ, JB or Jc virgin males. **B)** Refractoriness: the fraction of females that did not re-mate after mating first with FEJ, JB or Jc males. The ability of the three types of males to keep mates non-receptive was being tested. Error bars represent 95% confidence intervals around the mean of four replicate populations and therefore can be used for visual hypothesis testing.

#### B) Ability to re-mate with an already mated female

When the second mate was the experimental male, the ANOVA on mean refractory period showed no significant effect of male type (FEJ, JB or Jc) (*F*_*2,6*_=0.068*, P*=0.934) (Fig. 4A). However, there was a significant main effect of male type on refractoriness (*F*_*2,6*_=43.631*, P<=*0.001) (Fig. 4B). Pairwise comparisons revealed that the fraction of females which did not mate a second time was significantly less when the second mates were JB compared to FEJ and Jc males, indicating that the large JB males were more successful at re-mating with a non-virgin female, compared to the smaller FEJ or Jc males. FEJ males were the least successful at obtaining mating while the Jc male were intermediate between JB and FEJ (Fig 4B).

**Fig. 4.**
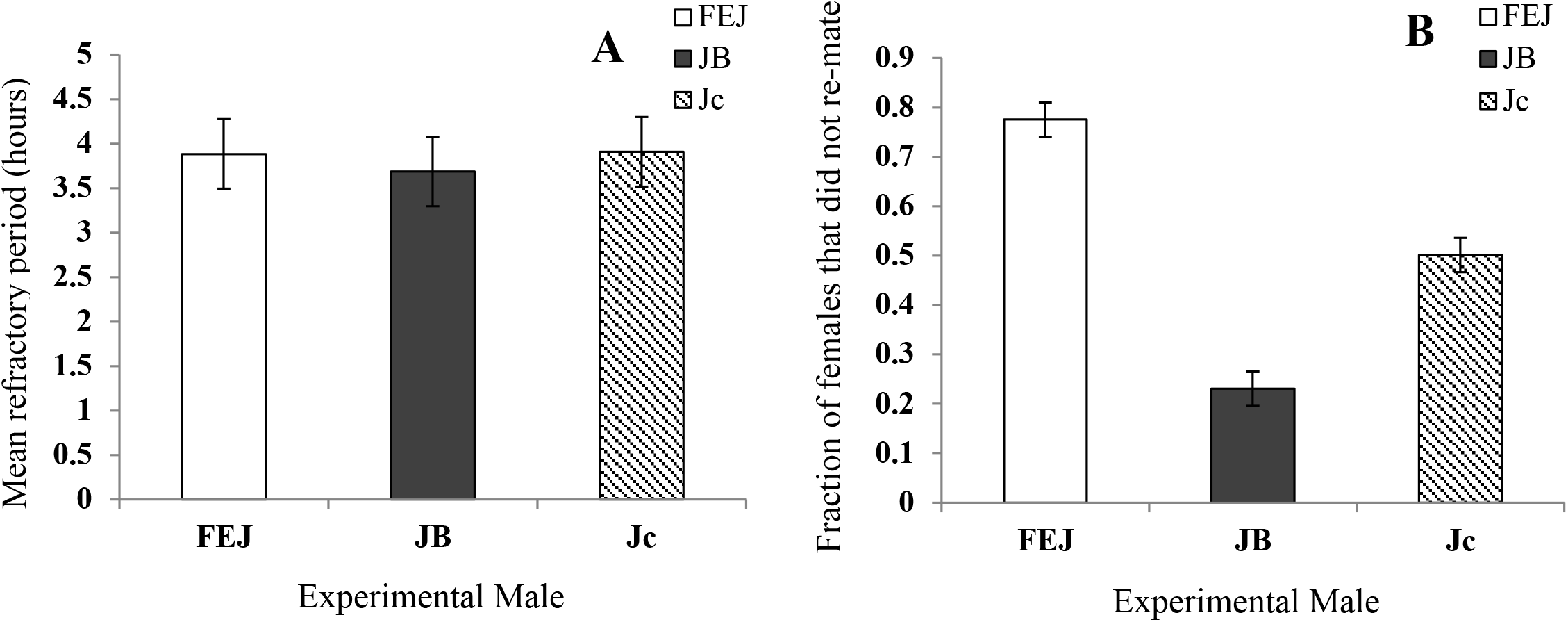
**A)** Mean refractory period: time till successful mating from introduction of second mates (FEJ, JB or Jc virgin males). **B)** Refractoriness: the fraction of females that did not re-mate. The ability of the three types of males to mate with an already mated female was being tested. Error bars represent 95% confidence intervals around the mean of four replicate populations and therefore can be used for visual hypothesis testing.

## Discussion

### Effect of body size reduction on post mating harm and manipulation caused by males

Our results from the post mating female mortality experiments indicate that a reduction in body size of males reduces the amount of post mating harm to females (more females died when mated with JB males than with Jc or FEJ males, on average) consistent with earlier work (Pitnick and Garcia-Gonzales 2002, M Ghosh and Joshi 2012). Mate harm in *Drosophila* is caused mainly by Acps transferred during copulation that perform a number of functions directed towards improving male fitness (Chapman et al. 2003), often at the expense of female fitness (Chapman et al. 1995). The complement of Acps that is produced by a *Drosophila* male is large, with redundancies, possibly as a response to resistance that females evolve against these proteins (Holland and Rice 1998, Sirot et al. 2015). Therefore, a change in the complement of Acps, the amount transferred, or both can determine the success of the manipulation and harm induced by a male. In our work, we assume Jc males to be producing the same set of Acps as JB males since they are genetically the same and speculate that the lower post-mating mortality induced by Jc than JB males is probably a result of reduced amount of Acps transferred due to their smaller body size.

Further evidence that Jc and JBs differ in post mating female manipulation is provided by the first of the refractory period assays (Fig 3 A, B) where the first mates were the experimental males. JB males induced a greater refractory period in their mates and succeeded more in preventing them from mating with a second male, than Jc or FEJ males.

In the second refractory period assay, the experimental males were the second mates, which showed that JB males were better at coercing/ successfully courting a non-virgin female into mating, than Jc or FEJ males (Fig 4B). In both refractory period assays however, Jc male performance was better than FEJ males’, unlike the results from the mortality assay. Taken together, while harm induced by males seemed to be entirely dependent on the size of the male, overall manipulative ability seemed to be affected both by their body size and genetic background. These results confirm that a reduction in the level of inter-locus sexual conflict in the FEJ population is, at least partly, a consequence of body size reduction. Perhaps, these traits are too expensive for FEJ males to exhibit.

### Effect of body size reduction on female defense

The post mating mortality experiments show that resistance to mating stress in females is unaffected by change in body size, as Jc and JB females have similar post mating mortality, both significantly lower than that of the FEJ females, irrespective of the type of mate to which they are exposed (Figs. 1, 2(A)). Since FEJ females are genetically different from JB and Jc females, our results suggest that female defense is affected by genetic background and not by body size. This result is consistent with a previous study on female body size and resistance to mate harm where large and small females were exposed to mates for a similar duration as ours (Long et al. 2009), although the female size variation used was much less than ours, as these females were taken from within a population and culturing condition.

It is important to note that most of the differences seen in the general trends of male induced female mortality were driven by the very high mortality rates of FEJ females paired with JB males indicating that, FEJ females are somehow “weak” compared to JBs. We found that a single generation reduction in body size resulted in males being less harmful towards females without females becoming more susceptible. The defence mechanisms to cope with biochemical aspects of mating stress were probably retained in Jc females but compromised in FEJ females. FEJ females, which have been co-evolving with smaller, less harmful males for many generations, have likely been selected to forgo any defenses against male harm, explaining the genetically “weaker” condition of FEJs. This would be a valuable resource saving for FEJ females, possibly allowing improved relative investment in gamete production. FEJ females show increased output per unit dry weight than JB (Prasad 2004, Ghosh-Modak 2009), consistent with this argument.

### Effect of larval crowding

We must consider the manner of production of Jc males here, i.e., larval crowding, which has also been used in many other studies (Jagadeeshan et al. 2015, Shenoi et al. 2016, Morrimoto et al. 2016, Wigby et al. 2016), as our results might be confounded by the effects of culturing condition and may not necessarily be due only to body size differences. In the post mating mortality assay, stress associated with courtship and mating, in addition to Acps transferred during mating, can bring about higher mortality rates in females.

Low food levels in the larval stage, i.e., high density conditions, might trigger a higher mating effort in the adult males due to the anticipated presence of a large number of males and, therefore, greater competition (Gage 1995, Wigby et al. 2016). In our experiments, it is not possible to distinguish between the effects of low body size per se, and contribution of larval crowding on the adult male’s behaviour. However, Edward and Chapman (2012) have demonstrated that crowding of larvae, by itself, does not affect mating related behaviour, so long as body size of the adults is not significantly affected. Where researchers have reported an effect of larval crowding, body size has also been significantly reduced (Jagadeeshan et al. 2015, Shenoi et al. 2016, Wigby et al. 2016).

Moreover, our assumption that these mortality differences are primarily a product of post-mating manipulative harm is validated by our observations of total exposure to copulation (reported in supplementary material), where both total duration and number of matings were estimated. These experiments were performed under identical conditions as the post-mating mortality assay and did not exhibit any trends that explain the mortality rate differences (Supplementary Material, Figs. S1, S2).

However, given that larval crowding can affect copulation duration and might also affect courtship through body size differences (Gage 1995, Wigby et al. 2016), our results on overall mating exposure might appear to be in contradiction of both other work on *D. melanogaster* (Jagadeeshan et al. 2015, Morrimoto et al. 2016, Shenoi et al. 2016, Wigby et al. 2016), as well as results from our own refractory period assays, which have shown various mating related traits of males to be affected by size reduction via larval crowding. Given the differences in mating ability with non-virgins (Fig 4B) and ability to keep mates non-receptive (Fig 3 A, B), it is surprising that in our populations, size reduction did not translate into a difference in mating number or total copulation duration, as a function of body size, when observed over 50 hours (see Supplementary Material, Figs S1-S4). We believe that the absence of competitors for that long a duration (Jagadeeshan et al. 2015, Morrimoto et al. 2016, Wigby et al. 2016 used much shorter exposure times), could have resulted in disinterest in mates over time, even in reproductively more active males (like JB), thus, somewhat equalizing the total mating exposure. Although our data show various trends, they are insufficient to explain the mortality rate differences seen among the FEJ, JB and Jc females.

### Conclusion

Taken together, our data confirm that in *D. melanogaster,* male ability to manipulate a female’s behaviour and physiology (post mating) to its advantage is significantly compromised by reduction in body size. A female’s resistance to this manipulation, however, is not affected by a reduction in body size per se, but probably via co-evolutionary processes. The role of sexual conflict in mediating speciation has been discussed (Ritchie 2007) and demonstrated in *D. melanogaster* (Syed et al. 2017). Since these populations experience some degree of reproductive isolation from their ancestral controls, likely through evolution of reduced inter-locus sexual conflict (M Ghosh and Joshi 2012), this is further evidence for the role of sexually antagonistic traits in mediating speciation. However, this work directly implicates long-term selection for rapid development, and not direct selection for sexually antagonistic traits in bringing about incipient reproductive isolation in *D. melanogaster*.

## References

Arnqvist G, Rowe L. Sexual conflict. Princeton university press; 2005.

Bangham J, Chapman T, Partridge L. Effects of body size, accessory gland and testis size on pre- and postcopulatory success in *Drosophila melanogaster*. Anim Behav. 2002;64: 915–921.

Bateman AJ. Intra-sexual selection in *Drosophila*. Heredity. 1948;2: 349–368.

Bonduriansky R, Maklakov A, Zajitschek F, Brooks R. Sexual selection, sexual conflict and the evolution of ageing and life span. Func Ecol. 2008;22: 443–453

Chapman T, Arnqvist G, Bangham J, Rowe L. Sexual conflict. Trends Ecol Evol. 2003;18: 41–47.

Chapman T, Liddle LF, Kalb JM, Wolfner MF, Partridge L. Cost of mating in *Drosophila melanogaster* females is mediated by male accessory gland products. Nature. 1995;373: 241–244.

Chippindale AK, Alipaz JA, Chen H-W, Rose MR. Experimental evolution of accelerated development in *Drosophila*. 1. Developmental speed and larval survival. Evolution. 1997;51: 1536–1551.

Edward DA, Chapman T. Sex-specific effects of developmental environment on reproductive trait expression in *Drosophila melanogaster*: Environmental variation and reproduction. Ecol Evol. 2012;2: 1362–1370.

Friberg U, Arnqvist G. Fitness effects of female mate choice: preferred males are detrimental for *Drosophila melanogaster* females. J Evol Biol. 2003;16: 797–811.

Gage MJG. Continuous variation in reproductive strategy as an adaptive response to population density in the moth *Plodia interpunctella*. Proc B Biol Sci. 1995;261: 25–30.

Holland B, Rice WR. Perspective: Chase-away sexual selection: Antagonistic seduction versus resistance. Evolution. 1998;52: 1–7.

Jagadeeshan S, Shah U, Chakrabarti D, Singh RS. Female choice or male sex drive? The advantages of male body size during mating in *Drosophila melanogaster*. PLoS One. 2015;10: e0144672.

Lefranc A, Bundgaard J. The influence of male and female body size on copulation duration and fecundity in *Drosophila melanogaster*. Hereditas. 2000;132: 243–247.

Long TAF, Pischedda A, Stewart AD, Rice WR. A cost of sexual attractiveness to high-fitness females. PLoS Biol. 2009;7: e1000254.

Lüpold S, Manier MK, Ala-Honkola O, Bélote JM, Pitnick S. Male *Drosophila melanogaster* adjust ejaculate size based on female mating status, fecundity and age. Behav Ecol. 2010;22: 184–191.

M Ghosh S, Joshi A. Evolution of reproductive isolation as a by-product of divergent life-history evolution in laboratory populations of *Drosophila melanogaster*. Ecol Evol. 2012;2: 3214–3226.

Markow TA, Ricker JP. Male size, developmental stability, and mating success in natural populations of three *Drosophila* species. Heredity. 1992;69: 122–127.

Markow TA. Reproductive behavior of *Drosophila melanogaster* and *D. nigrospiracula* in the field and in the laboratory. J Comp Psychol. 1988;102: 169.

Modak SG, Satish KM, Mohan J, Dey S, Raghavendra N, Shakarad M, et al. A possible tradeoff between developmental rate and pathogen resistance in *Drosophila melanogaster*. J Genet. 2009;88: 253–256.

Morimoto J, Pizzari T, Wigby S. Developmental environment effects on sexual selection in male and female *Drosophila melanogaster*. PLoS One. 2016;11: e0154468.

Nandy B, Joshi A, AliZS, Sen S, Prasad NG. Degree of adaptive male mate choice is positively correlated with female quality variance. Sci Rep. 2012;2: 447.

Nunney L. The response to selection for fast larval development in *Drosophila melanogaster* and its effect on adult weight: An example of a fitness trade-off. Evolution. 1996;50: 1193–1204.

Parker GA. Sexual selection and sexual conflict. Blum MS & Blum NA, (Eds.), Sexual selection and reproductive competition in insects. NY: Academic Press. 1979. 123–166.

Partridge L, Farquhar M. Lifetime mating success of male fruitflies (*Drosophila melanogaster*) is related to their size. Anim Behav. 1983;31: 871–877.

Partridge L, Ewing A, Chandler A. Male size and mating success in *Drosophila melanogaster*: the roles of male and female behaviour. Anim Behav. 1987a;35: 555–562.

Partridge L, Hoffmann A, Jones JS. Male size and mating success in *Drosophila melanogaster* and *D. pseudoobscura* under field conditions. Anim Behav. 1987b;35: 468–476.

Partridge L, Fowler K. Non-mating costs of exposure to males in female *Drosophila melanogaster*. J In Physiol. 1990;36: 419–425.

Pitnick S, García-González F. Harm to females increases with male body size in *Drosophila melanogaster*. Proc Biol Sci. 2002;269: 1821–1828.

Prasad NG, Shakarad M, Anitha D, Rajamani M, Joshi A. Correlated responses to selection for faster development and early reproduction in *Drosophila*: the evolution of larval traits. Evolution. 2001;55: 1363–1372.

Prasad NG, Shakarad M, Gohil VM, Sheeba V, Rajamani M, Joshi A. Evolution of reduced pre-adult viability and larval growth rate in laboratory populations of *Drosophila melanogaster* selected for shorter development time. Genet Res. 2000;76: 249–259.

Prasad NG. Life-history evolution in laboratory populations of *Drosophila melanogaster* subjected to selection for faster development and early reproduction. Jawaharlal Nehru Centre for Advanced Scientific Research. 2003.

Rice WR. Dangerous liaisons. Proc Natl Acad Sci U S A. 2000;97: 12953–12955.

Ritchie MG. Sexual Selection and Speciation. Annu Rev Ecol Evol Syst. 2007;38: 79–102.

Trivers R. Parental investment and sexual selection. B Cambell (Ed), Sexual selection and the descent of man. Aldine DeGruyter. NY, USA. 1972; 136–179.

Roff, DA. Life history evolution. Sinauer Associates, Inc. Sunderland, MA, USA. 2002.

Satish KM. Reverse evolution and gene expression studies on populations of *Drosophila melanogaster* selected for rapid pre-adult development and early reproduction. Jawaharlal Nehru Centre for Advanced Scientific Research. 2010.

Shenoi VN, Banerjee SM, Guruswamy B, Sen S, Ali SZ, Prasad NG. *Drosophila melanogaster* males evolve increased courtship as a correlated response to larval crowding. Anim Behav. 2016;120: 183–193.

Sirot LK, Wong A, Chapman T, Wolfner MF. Sexual conflict and seminal fluid proteins: a dynamic landscape of sexual interactions. Cold Spring Harb Perspect Biol. 2014;7: a017533.

StatSoft, Inc. STATISTICA for Windows. Tulsa, OK, USA. 1995

Syed ZA, Chatterjee M, Samant, MA, Prasad NG. Reproductive isolation through experimental manipulation of sexually antagonistic co-evolution in *Drosophila melanogaster*. Sci Rep. 2017;7: 3330.

Wigby S, Perry JC, Kim Y-H, Sirot LK. Developmental environment mediates male seminal protein investment in *Drosophila melanogaster*. Func Ecol. 2016;30: 410–419.

Zwaan B, Bijlsma R, Hoekstra RF. Artificial selection for development time in *Drosophila melanogaster* in relation to the evolution of ageing: Direct and correlated responses. Evolution. 1995;49: 635–648.

