## Supplementary Material for "Evolution of lower levels of inter-locus sexual conflict in *D. melanogaster* populations under strong selection for rapid development"

### Methods

*Mating exposure in 50 hours*

Since the JB males are larger than the Jc or FEJ males, the increased mortality of females mated to JB males could be due to increased number of matings when housed with JB males rather than an intrinsically higher mate-harming ability of JB males. We therefore checked for differences in number of matings in the nine different combinations of male and female type. Similar to the set up described for estimating post-mating female mortality, single pairs were set up when the virgin flies were 3-4 days old, and the number of successful matings in 50 hours recorded for each male-female combination. Ten vials were set up per combination of male-female type and all vials were under continuous observation in a well illuminated environment without disturbance and a constant temperature of $25^{o}C \pm 1^{o}C$ for a duration of 50 hours from setup. A mating was considered successful only when the mating pair was observed to be in copula for at least three minutes. We noted the duration of each mating and calculated the total mating exposure of each female since that could also significantly contribute to the stress experienced by females.

### Results

*Mating exposure in 50 hours*

ANOVA results showed significant effects of female type, male type and male type × female type interaction for mean number of matings in 50 hours of exposure (Table S1). JB females with JB males showed maximum number of matings, significantly higher than the other combinations, as revealed by pairwise comparisons (Fig. S1). This was mirrored when the total duration of mating (total exposure to mating) was analyzed with the main effects of female type, male type and male type × female type interactions being significant (Table S2) (Fig. S2, S4). Trends also revealed that Jc females, in general, were mated with less than JB females, although this was significantly different only when total mating duration was checked (Fig. S2) (Table S2). This might be indicative of male mate preference, where larger females have been shown to be preferred (Anderson 1994, Byrne and Rice 2006), or better ability of the Jc females to avoid being force-mated with. The higher activity levels of Jc females is a more likely cause, since a single female is available for the male, and therefore, choice cannot be exercised in the absence of another female. Jc males were significantly more successful at mating compared to FEJ males in terms of both the mean number of matings achieved (Fig. S3 (B)) and total duration of mating exposure (Fig. S4 (B)) and did not significantly differ from JB males in either measure of male mating success. This can be indicative of better quality of Jc males compared to FEJ males, even though they are of equally small size.

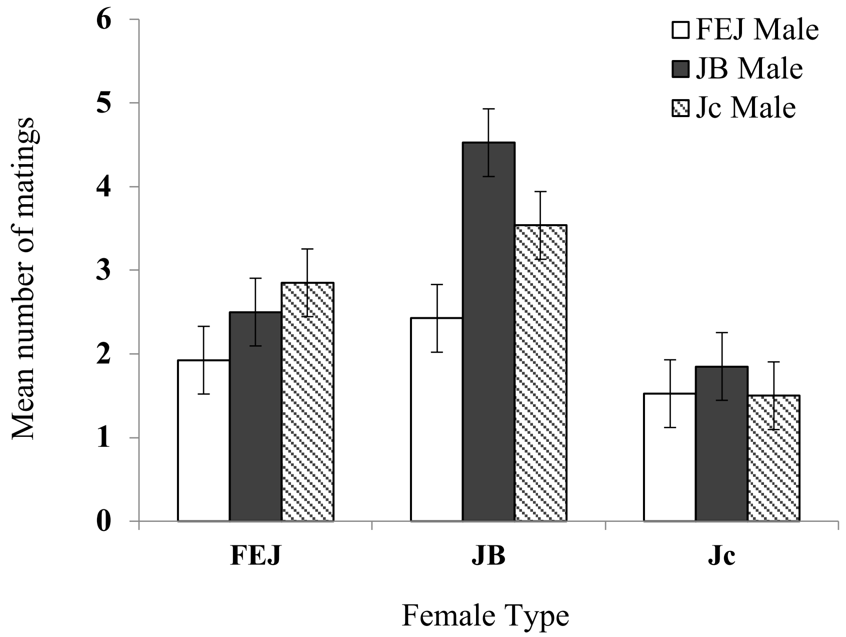


Fig. S1 Mean number of matings as recorded in 50 hours of females being housed with FEJ, JB or Jc males. Error bars are 95% confidence intervals and can therefore be used for visual hypothesis testing.

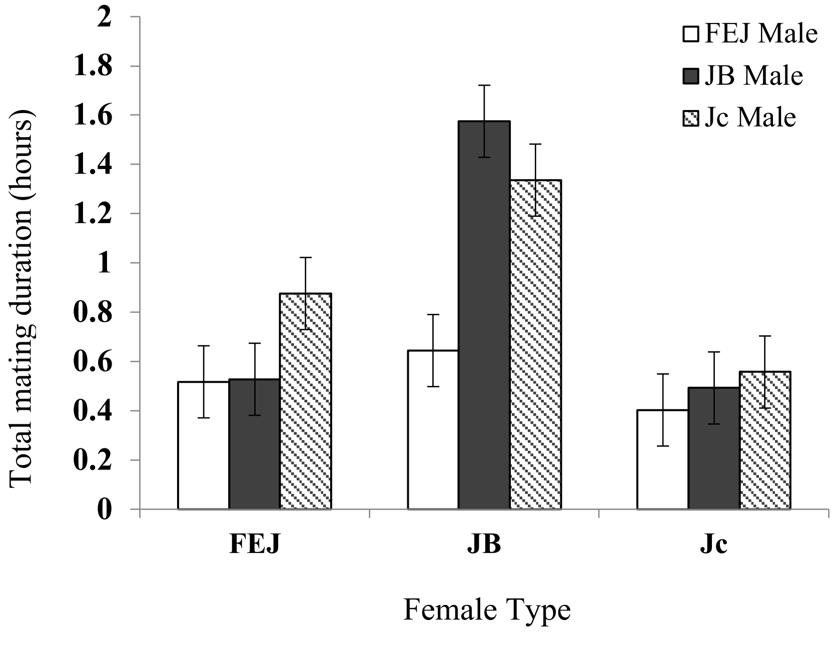


Fig. S2 Total exposure to mating as experienced by females when housed for 50 hours with FEJ, JB or Jc males. Error bars are 95% confidence intervals, facilitating visual hypothesis testing.

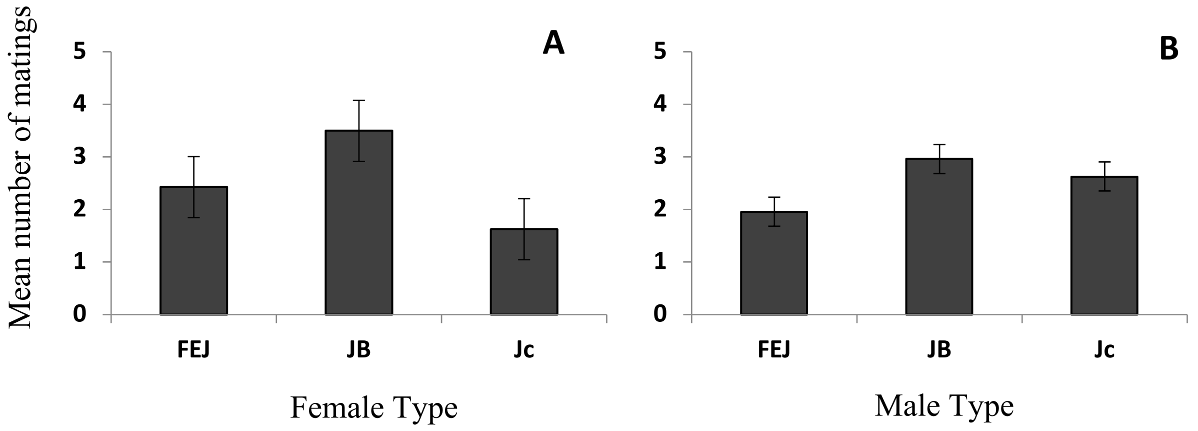


Fig. S3 Mean number of matings in 50 hours of exposure showing (A) main effect of female type, and (B) main effect of male type. Error bars are 95% confidence intervals and can therefore be used for visual hypothesis testing.

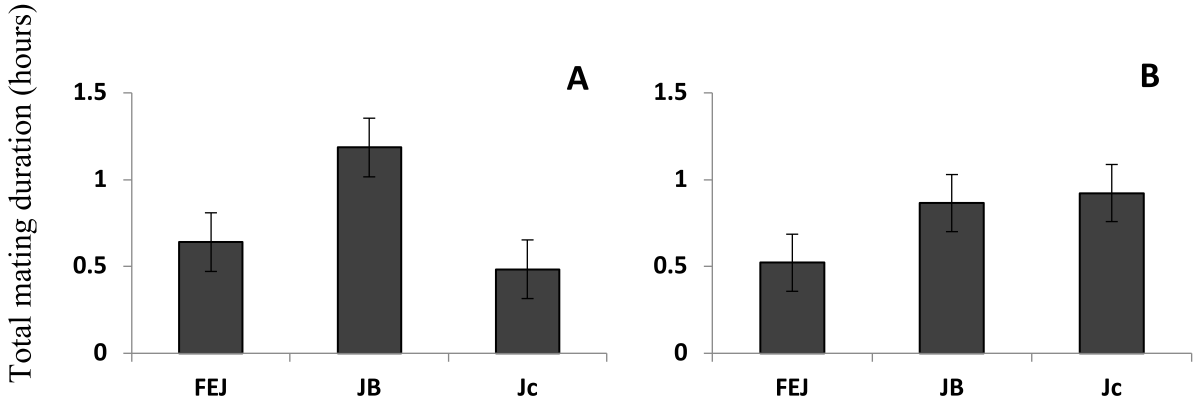


Fig. S4 Total exposure to mating in 50 hours showing (A) main effect of female type, and (B) main effect of male type. Error bars are 95% confidence intervals and can therefore be used for visual hypothesis testing.

| Effect | *df* | MS | *F* | *P* |
|---|---|---|---|---|
| Female | 2 | 10.567 | 17.70 | 0.0030 |
| Male | 2 | 3.1161 | 23.06 | 0.0015 |
| Female × Male | 4 | 1.1619 | 4.004 | 0.0273 |

Table S1 Results from ANOVA performed on mean number of matings in 50 hours, with male and female type as fixed factors. In this design random factors and interactions are not tested for significance and have been omitted for brevity.

| Effect | *df* | MS | *F* | *P* |
|---|---|---|---|---|
| Female | 2 | 1.625 | 32.15 | 0.0006 |
| Male | 2 | 0.565 | 11.80 | 0.0083 |
| Female × Male | 4 | 0.280 | 7.385 | 0.0030 |

Table S2 Results from the ANOVA performed on total duration of matings exposure in 50 hours, with male and female type as fixed factors. In this design random factors and interactions are not tested for significance and have been omitted for brevity.
